# Spatial localization of recent ancestors for admixed individuals

**DOI:** 10.1101/004713

**Authors:** Wen-Yun Yang, Alexander Platt, Charleston Wen-Kai Chiang, Eleazar Eskin, John Novembre, Bogdan Pasaniuc

## Abstract

Ancestry analysis from genetic data plays a critical role in studies of human disease and evolution. Recent work has introduced explicit models for the geographic distribution of genetic variation and has shown that such explicit models yield superior accuracy in ancestry inference over non-model-based methods. Here we extend such work to introduce a method that models admixture between ancestors from multiple sources across a geographic continuum. We devise efficient algorithms based on hidden Markov models to localize on a map the recent ancestors (e.g. grandparents) of admixed individuals, joint with assigning ancestry at each locus in the genome. We validate our methods using empirical data from individuals with mixed European ancestry from the POPRES study and show that our approach is able to localize their recent ancestors within an average of 470Km of the reported locations of their grandparents. Furthermore, simulations from real POPRES genotype data show that our method attains high accuracy in localizing recent ancestors of admixed individuals in Europe (an average of 550Km from their true location for localization of 2 ancestries in Europe, 4 generations ago). We explore the limits of ancestry localization under our approach and find that performance decreases as the number of distinct ancestries and generations since admixture increases. Finally, we build a map of expected localization accuracy across admixed individuals according to the location of origin within Europe of their ancestors.

**Author Summary:** Inferring ancestry from genetic data forms a fundamental problem with applications ranging from localizing disease genes to inference of human history. Recent approaches have introduced models of genetic variation as a function of geography and have shown that such models yield high accuracies in ancestry inference from genetic data. In this work we propose methods for modeling the mixing of genetic data from different sources (i.e. admixture process) in a genetic-geographic continuum and show that using these methods we can accurately infer the ancestry of the recent ancestors (e.g. grandparents) from genetic data.

## Introduction

Inference of ancestry from genetic data is a critical aspect of genetic studies with applications ranging from mapping genes to diseases to the inference of population history [29, 33, 32, 9, 31, 20]. Although many initial large scale genetic association studies have focused primarily on homogeneous populations, increasingly studies are addressing samples in which individuals have more complex backgrounds, including admixed ancestry (i.e. emerging from the mixing of genetically diverged ancestors) [13, 12, 38, 7, 18, 26]. Such studies depend crucially on accurate and unbiased ancestry inference both at a genome-wide level as well as at each locus in the genome [33, 22].

Traditional ancestry inference from genetic data has been focused on modeling populations as discrete units. As a result, traditional genome-wide ancestry inference estimates the proportion of sites in the genome coming from a set of source populations (continental or subcontinental), and locus-specific inference aims to assign each allele in the genome to one of the considered populations [10, 30, 2, 1, 28, 22, 35]. More recent approaches model population structure in a geographic continuum capitalizing on the correlation of genetics and geography expected in isolation by distance models [27, 39, 3, 37] and observed in many organisms [41, 15]. This has most often been performed through principal components analysis (PCA) [6, 37, 20, 34, 27, 24, 23, 40, 17], a general procedure for reducing the dimensionality of the data, with more recent approaches focusing on explicit modeling of the relationship between patterns of genetic variation and geography [39, 3]. These approaches typically assume that an individual’s genotype is drawn from the genetic variation present at a single geographic location. This assumption is clearly violated when individuals have ancestors from multiple geographic regions, as occurs with recently admixed populations in the Americas (such as African-Americans) and more generally, individuals that have ancestry from multiple regions within the same continent (e.g. individuals with recent ancestors from multiple regions of Europe). Recent approaches have circumvented this issue by first inferring segments of different continental ancestry (i.e. locus-specific) followed by independent application of PCA only on segments of specific continental ancestry (e.g. European segments) [14]. A critical component of this approach is the performance of locus-specific ancestry inference which has been shown to attain high accuracy for continental ancestries but less accurate in inferring sub-continental ancestry (e.g. country of origin)[28, 2, 16]. A different approach is to model admixed individuals directly as function of geography. For example, the SPA [39] method allows for the limited scenario where an individual is a descendant of parents who are not themselves admixed, but are from different locations in Europe.

In this work we introduce models of admixture across varying number of generations and ancestries in a geographic continuum. We model admixed genomes as having recent ancestors from several locations on a genetic-geographical map. We perform ancestry inference by simultaneously localizing on the map the recent ancestors of an admixed individual and by partitioning the admixed genome into segments inherited from the same ancestor (i.e. locus-specific ancestry). We take advantage of the simple observation that if one allele is inherited from a specific ancestor, then most likely the neighboring alleles are also inherited from the same ancestor. Specifically, we use a model-based framework for genetic variation in the geographical continuum [39] and employ a hidden Markov modeling of the admixture process [25, 11]. We develop efficient optimization algorithms that allow us to accurately predict the geographic location of the recent ancestors of an admixed individual in conjunction with locus-specific ancestry inference. The results allow the localization on a geographical map of each allele in recently admixed individuals.

We use empirical genotype data from the Population Reference Sample (POPRES) project [19] to validate our approach. The POPRES project has genotyped more than 3,000 individuals with ancestry distributed throughout Europe and has recorded the self-reported ancestry (typically at the level of country) for both individuals and their parents/grandparents. We use the POPRES individuals with homogeneous ancestry (i.e. all grandparents having the same reported ancestry) to infer patterns of variation across geography in Europe [39] and employ our method to localize the recent ancestors of individuals with self-reported admixed ancestry (i.e. grandparents with multiple ancestries in Europe). Our method is able to localize the grandparents of the admixed individuals in POPRES data within an average of 470Km of their reported ancestry. The accuracy is dependent on the specific ancestries and ranges from 305Km for individuals with Swiss and French ancestry to 701Km for those with Spanish and Portuguese ancestry. We use simulations from POPRES genotype data to show that the localization accuracy within Europe decreases with increased number of ancestors and with the number of generations since the admixture. We also show that inference accuracy (at the genome-wide and locus-specific level) increases as distance among ancestors increases. Finally, we provide an analysis of ancestry localization error across all pairs of countries in Europe as resource for community interested in subcontinental ancestry in Europe. A software package implementing our methods is freely available at *http://bogdan.bioinformatics.ucla.edu/software/*.

## Results

### Overview of spatial localization for admixed individuals

In this work we consider models of ancestry for admixed individuals in a geographical continuum. We view the mixed ancestry genome as being generated from several geographical locations on a map, corresponding to the locations of their recent ancestors (see Figure 1). For example, consider the case of an individual with one maternal grand-parent from Italy and the other one from Great Britain (see Figure 1a). The maternal copy of its genome will be composed of segments originating from the two locations in Europe (see Figure 1b). Each position in the genome has its own function that describes the population allele frequencies at that site as a function of geography. The approach we take follows SPA[39] and assumes these functions take on logistic gradient shapes. Some variants may have steep gradients (i.e. frequencies that change drastically with location) while other variants may not vary at all with geography (see Figure 1c). Although these types of functions clearly do not explain all correlation between genetics and geography, it has been previously shown that such simple functions carry sufficient information to be very informative of ancestry status across individuals [39, 3] and lead to likelihood functions that are simple to optimize. Other types of functions such as linear [3] or more complex could also be employed in our framework.

**Figure 1:**
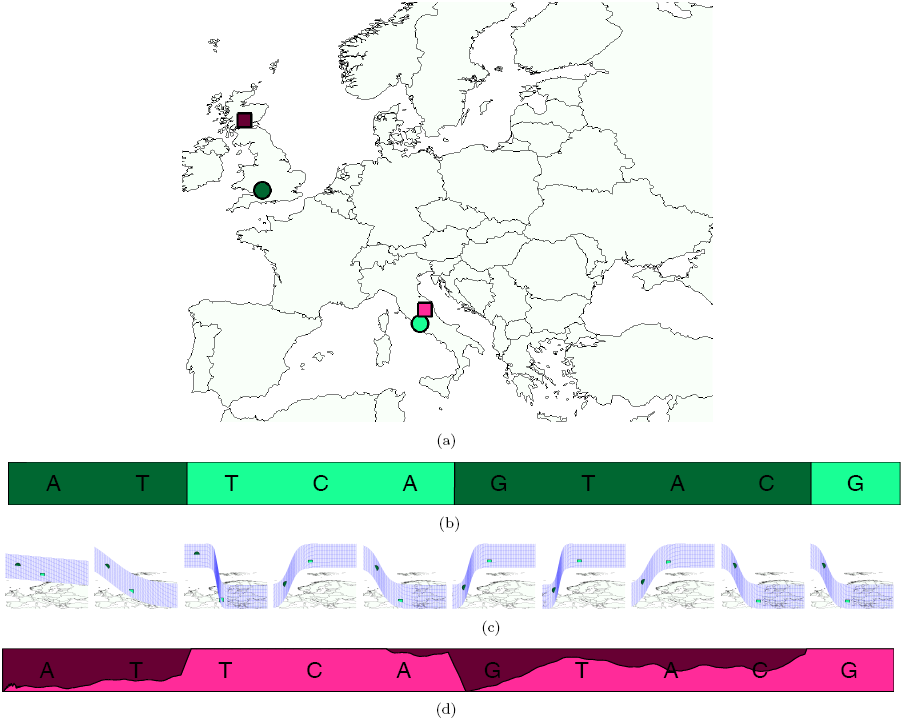
SPAMIX model for admixed individuals. (a) Example of haploid individual with two ancestry locations in Europe (circles denote the true ancestry locations). (b) The admixture process induces segments of different ancestry backgrounds. (c) SPAMIX uses logistic gradients to describe allele frequencies as a function of geographic map to instantiate an admixture HMM for each pair of locations on a map. Each location on the map is associated to a particular allele frequency at all sites in the genome. (d) SPAMIX finds the location of ancestors on a map (denoted by squares in subfigure (a)) and the locus-specific ancestry at each site in the genome by maximizing the likelihood of genotype data.

Having estimated gradient functions at each site in the genome, we extend standard hidden Markov models (HMM) for admixture to incorporate variation at each position on the map by allowing the emission probabilities to vary according to these gradients (see Methods). We perform inference in this model to find the ancestor locations on the map that maximize the likelihood of the observed genome (Figure 1a). After finding the location of the recent ancestors, we assign each allele in the mixed genome to one of the ancestor locations. This provides a locus-specific ancestry call across the genome. Figure 1d shows an output of our method (SPAMIX) with locations in the admixed genome being labeled according to the inferred ancestral location on the map.

Although we presented a simple example of our framework with two ancestors on the map, the model flexibly handles diploid data with arbitrary number of generations and ancestors localized on the map (e.g. diploid genome with 4 ancestors localized on the map 2 generations ago, diploid genome with 8 ancestor locations 3 generations ago) (see Methods). As the number of generations since admixture increases, the total number of ancestors to localize increases dramatically, making the inference problem very challenging. To account for this effect we limit the number of different locations for the recent ancestors for the maternal (paternal) haplotypes to a fixed constant (e.g. *M*(*N*), see Methods) with varying amount of contributions to the admixture process (see Methods). We devise efficient algorithms to jointly optimize the locations of the ancestors as well as the proportion they contribute to the genome of the admixed individual (see Methods). For example, in the case of 3 generations ago, one ancestry location may contribute 1/8^*th*^ to the admixed genome if only one ancestor comes from that location and may contribute 1/2 to the admixture process if half the ancestors come from that specific location. Finally, we note that the diploid model is symmetric making *M* and *N* interchangeable.

### Performance of continuous ancestry inference in simulations

We investigated the performance of our model through simulations from the Population Reference Sample (POPRES) data [19]. The POPRES data measures genome-wide genetic variation in a large number of individuals with ancestries across Europe (with a larger proportion of individuals with ancestry from the central and western Europe). For each individual, the self-reported ancestry (typically at the level of country) of parents and grandparents was recorded. To produce a large number of admixed individuals on which to test the data, we first randomly selected individuals with homogenous ancestry (i.e. all 4 grandparents from the same country of origin) from various areas in Europe to serve as ”ancestor” genomes. We used them to simulate an admixed individual and attempted to recover the original ancestral locations using the simulated genome and a set of logistic gradients inferred from the remaining unused individuals (see Methods). SPAMIX attains an ancestry localization accuracy (i.e. average distance between true and inferred locations of the recent ancestors) for individuals with two recent ancestors in Europe of 550 Km (see Table 2). There is a large variance (334Km) across different sets of ancestral pairs showing the high variability in performance across subjects. Contributing to this effect is the variable sampling density of the POPRES data (which is denser towards the center of Europe) and variability of goodness-of-fit of the logistic gradients across the map (e.g. poorer fit at range boundaries).

**Table 1.** Self-reported grandparental ancestry (location of origin) of the POPRES data individuals (1906 in total). For individuals with grandparental ancestry from 2 different countries, we also report the number of individuals with 2 grandparents from one location and 2 from the other (2/2), versus individuals with 3 grandparents from one country (3/1).

|  | Number of Different Ancestries |  |  |  |  |  |
| --- | --- | --- | --- | --- | --- | --- |
|  | 1 | 2 |  |  | 3 | 4 |
|  |  | (2/2) | (3/1) | Total |  |  |
| Number of Individuals | 1385 | 261 | 153 | 414 | 54 | 2 |
| Percentage out of total | 74.7% | 14.1% | 8.2% | 22.3% | 2.9% | 0.1% |

**Table 2.** Average distance between inferred and true ancestry locations in simulated admixed individuals from POPRES data. Simulations assume 4 generations in the mixture process. Naive model denotes the extension of SPA that ignores admixture-LD. SPAMIX (logistic) represents simulation results starting from haplotypes generated at a location on a map using a Bernoulli sampling from the logistic gradients (see Methods). Parenthesis denote the standard deviation while standard error of the mean is computed as standard deviation divided by square of number of simulations in each category.

| No. of ancestries | 1 | 2 | 3 | 4 |
| --- | --- | --- | --- | --- |
| Naive model | $443 \pm 4(265)$ | $880 \pm 5(491)$ | $898 \pm 10(530)$ | $880 \pm 9(578)$ |
| SPAMIX haploid model | $458 \pm 4(273)$ | $557 \pm 4(334)$ | $620 \pm 7(392)$ | $665 \pm 7(449)$ |
| SPAMIX diploid model | $443 \pm 4(265)$ | $550 \pm 4(326)$ | $591 \pm 7(367)$ | $639 \pm 7(423)$ |
| SPAMIX (logistic) | $75 \pm 1(41)$ | $236 \pm 5(131)$ | $363 \pm 6(215)$ | $419 \pm 6(247)$ |

To test how much is gained by explicit modeling of correlations among SNPs induced by segments of recent shared ancestry (admixture LD), we also inferred the recent ancestry location using a naïve model that assumes all SNPs to be independent (as in [39]). We observe a significant increase in the average distance between true and inferred locations in this naïve model (880 Km) versus the 550 Km for SPAMIX thus showing that modeling admixture LD significantly increases performance. We also quantified the effect of correlations among markers conditional on local ancestry (background LD) in our approach. Eliminating loci found in strong disequilibrium with each other (LD pruning) was observed to increase accuracy even though the model had less data to use (see Supplementary Tables 1 and 2); therefore, all results in the main text are obtained after LD pruning (r2*<*0.2, see Methods).

It is increasingly often the case that access to pedigree data allows haplotypes to be determined with high accuracy. Therefore, we quantified the gain in ancestry localization accuracy arising from having access to phased haplotype data (i.e. haploid data) as compared to unphased diploid data. Table 2 shows that accurate phasing significantly increases localization accuracy. For example, having access to perfect phasing allows for the inference of the 4 ancestral locations within 557Km of the simulated location where the diploid model attains an average of 639Km of its simulated locations.

An important parameter of our model is the number of generations since admixture; with more generations, more recombination events have the opportunity to shuffle ancestry across the genome thus reducing the average length of the ancestry segments. We observe a slight decrease in performance from 2 to 8 generations (548 to 562 Km) which we expect to continue as the number of generations increases (in the limit of extremely large number of generations, our model is equivalent to the naïve model that does not model admixture-LD) (see Table 3).

**Table 3.**
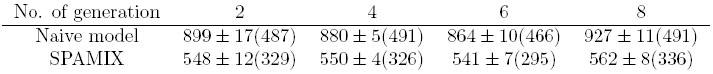
Average distance between inferred and true ancestry locations in simulated admixed individuals from POPRES data as function of number of generations in the mixture process. Two ancestral locations were assumed for this simulation. Parenthesis denote the standard deviation while standard error of the mean is computed as standard deviation divided by square of number of simulations in each category.

| No. of generation | 2 | 4 | 6 | 8 |
| --- | --- | --- | --- | --- |
| Naive model | $899 \pm 17(487)$ | $880 \pm 5(491)$ | $864 \pm 10(466)$ | $927 \pm 11(491)$ |
| SPAMIX | $548 \pm 12(329)$ | $550 \pm 4(326)$ | $541 \pm 7(295)$ | $562 \pm 8(336)$ |

Our framework models genetic variation as function of geography by imposing a logistic gradient to the generating functions (see Methods). That is, the frequency of a given variant is allowed to change in a given direction on a map only according to a parametrized logistic function. Although this approach has been shown to provide a good approximation of common variation leading to accurate ancestry inference, we hypothesize that the error in fitting logistic gradients to real data limits the method’s accuracy. To assess this scenario, instead of using real individual’s haplotype data, we simulated admixed haplotypes directly from the logistic gradients we inferred from POPRES data (see Methods). We observe a large increase in accuracy in this idealized scenario as compared to simulations from real haplotype data (e.g. 236 vs 550 Km for 2 ancestries 4 generations ago, Table 2), thus indicating that logistic gradients do not account for all the correlation between geography and genetic variation. This suggests that further work on functions linking geography to genetics within our framework may yield additional improvements (see Discussion).

We investigated the performance of our approach as we increase the number of ancestral locations (*M*/*N*, see Methods) to estimate for a given admixed individual. For a fixed number of generations (four), we varied the number of ancestry locations to estimate. The parental inference is different from two ancestry inference, as the parental inference assumes that one haplotype is from paternal ancestry and one from maternal ancestry. However, the two-ancestry inference assumes that both of the haplotypes are mosaic of two ancestries (*M* = *N* = 2). As expected, we observe decreases in performance as the number of ancestral locations increases. For example, the average prediction error increases from 550 for two ancestries to 639 Km for four ancestral locations (Table 2).

### Increased distance between ancestral locations improves performance

It is well known that accuracy of ancestry inference correlates with genetic distance between ancestral populations. Discrete local ancestry can be inferred with very high degree of accuracy in mixtures of highly diverged populations (e.g. African Americans) as compared to closely related ones (e.g. sub continental mixtures) [21, 4, 16, 28]. Since geography correlates with genetic distance, we hypothesized that the accuracy of continuous ancestry inference in recently admixed individuals also correlates with distance among ancestries on the map. Indeed, we observe that the relative prediction error (i.e. the difference between predicted and true locations normalized by the distance between the true ancestry locations, see Methods) decreases with the distance between ancestries in Europe (Figure 2A). For example, if the ancestries are 500 Km apart, we observe a relative prediction error of 0.75 as compared to 0.50 when the ancestries are located 2000 Km apart. Interestingly, when not normalizing for the distance between ancestries (Figure 2B), we observe that prediction error increases with increased distance. This shows that although the task of separating the ancestry locations becomes simpler, the localization accuracy becomes poorer (e.g. two ancestors located 500Km apart are localized within 450Km of their true locations, while two ancestors located 3000Km apart are localized within 1000Km of their true locations). This effect is presumably due to assignment errors in the local ancestry that have a much bigger impact if the ancestral locations are further apart. Although fewer local ancestry errors are being made with increased distance (see below), these errors have a stronger impact on the ancestral localization due to their higher distance to the true location.

**Figure 2:**
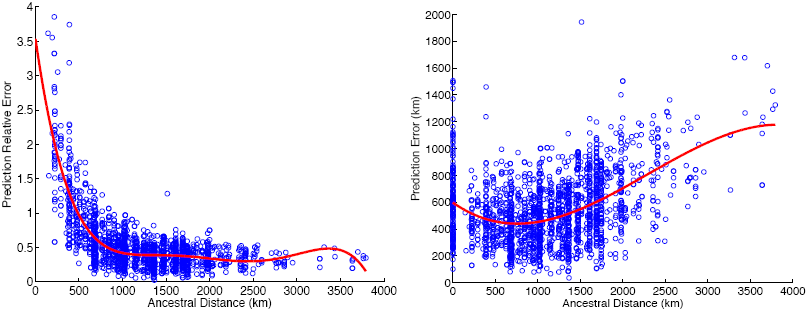
Ancestral location prediction error as a function of distance between ancestral locations in simulations over POPRES data. Left shows the prediction error normalized by the distance between the ancestral locations used in simulations and right plots the the prediction error. Simulations use the haploid model with 2 generations in the mixture.

### Inference of number of ancestors

In all simulations above we have assumed that the true number of different ancestry locations is known. We investigated whether our approach can also be used to predict the number of distinct ancestries on the map for a given genome. We used the standard Akaike information criterion (AIC)[5] that balances the goodness of fit with the number of parameters in the model (more ancestries to infer increases the number of parameters in our method). Starting from POPRES individuals with homogeneous ancestry we simulated admixed individuals with up to 4 ancestry locations 4 generations ago under the constraint that the ancestries are at least 600 kilometers apart. For each simulated admixed individual, we ran our method SPAMIX using *N* = 1, 2, 3, 4 ancestry locations and used the AIC to infer the number of ancestors. Figure 3 shows that this procedure will on average estimate the number of ancestries correctly, but the error rate is expected to be high for any single case of inference.

**Figure 3:**
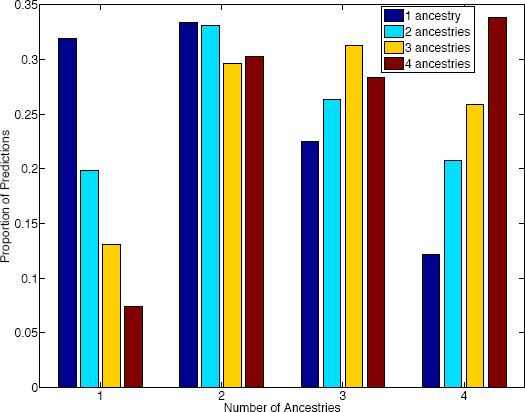
Inference of number of distinct ancestries using the Akaike information criterion (AIC). We simulated 1,000 admixed individuals with up to 4 distinct ancestry sources in Europe and used the AIC within the SPAMIX model to infer the number of ancestries. Figures show the proportion of inferred number of ancestries (y-axis) as function of number of simulated ancestries (x-axis) Although we observe a large variance in the number of predicted ancestries we note that the histogram is centered on the correct simulated number of ancestries thus suggesting that AIC could be employed to infer the number of distinct ancestors.

### Locus-specific inference

An advantage of our framework is that in addition to identifying the most likely locations of the recent ancestry of admixed individuals, it can also provide an assignment of each allele in the genome to each ancestry location. We observe that local ancestry prediction accuracy (i.e. the proportion of alleles assigned to the correct ancestry, see Methods) increases with the distance between ancestral locations (Figure 3) from 55% of loci assigned accurately for very closely related ancestries (less than 500Km apart) to more than 70% for ancestries 2500Km apart (Supplementary Figure 1). Similar to the ancestor localization, we observe that although the total number of assignment errors is reduced with increased distance, these errors have a bigger impact when averaging across all sites to compute the average allele localization error. Therefore, we observe that the average local ancestry prediction error is increased as distance between ancestral locations is increased.

### Map of accuracy across Europe

We also investigated the variance in performance according to the ancestor’s labeled origin (i.e. typically to level of country). Figure 4 shows the prediction error for admixed individuals with ancestry from pairs of origins in Europe. In general, we observe decreased performance for populations at the boundary of the European map (e.g., Portugal, Spain, Italy), and increased performance for subcontinental admixtures from populations located geographically in the center of Europe (e.g. France, Switzerland) (Supplementary Figures 2 and 3). This can be an effect of biased sampling in the POPRES data, that sampled more individuals from Europe center, but also can be an effect of having more information to localize individuals in SPAMIX. In general we observe a prediction accuracy ranging from 411 Km for admixtures from Spain and Italy to 641 Km for individuals with recent ancestors from Spain and the UK.

**Figure 4:**
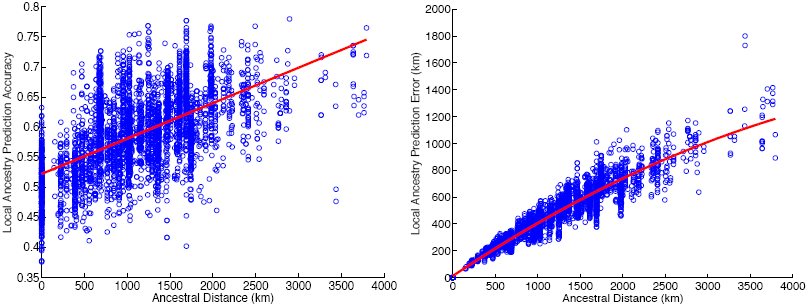
SPAMIX locus-specific ancestry prediction accuracy as function of distance between ancestral locations. Left displays the local ancestry prediction accuracy, defined as the percentage of all loci with correct assignment of ancestry. Right plot displays the average distance to true locations for each allele in the genome (local ancestry prediction error). Simulations use the haploid model with 2 generations in the mixture.

### Analysis real admixed individuals from POPRES data

Finally, we investigated whether high accuracies observed in simulations can also be attained in real data. Using SPAMIX, we localized the recent ancestry of all admixed European individuals from POPRES (see Methods). A total of 470 admixed individuals are analyzed using SPAMIX (see Table 1). As “ground truth” ancestral locations, we used the the center of the self-reported grandparent country of origin. Therefore, we assume the mixed individuals from POPRES have two to four ancestry locations to infer. Across all 470 individuals, we observe an average prediction error distance of 470Km which decreases to 426 if outlier individuals defined as those with prediction errors larger than 1, 000Km are removed (all such outlier individuals are reported in Supplementary Table 3). The error distance is lower than simulated experiments likely due to the large proportion of the admixed individuals of French and Swiss ancestries which can be accurately localized (average of 305Km). As above, we note that SPAMIX ancestor localization performance varies greatly across Europe with ancestors from pairs of countries localized at the boundary of European map being harder to localize (e.g. an average of 701Km for ancestor localization for Spanish/Italian mixed individuals).

## Discussion

We have introduced new models for predicting the geographical origins of multiple recent ancestors for individuals with recent mixed ancestry. Existing methods for ancestry inference in admixed populations either focus on discrete ancestry assignment or use locus-specific ancestry inference followed by PCA on subsets of the data. We introduce models that leverage the spatial structure of genetic variation using hidden Markov models for the admixture process to achieve high accuracy in localizing the recent ancestry of a given individual on a geographical map. Our proposed model can be viewed as a generalization of the parental localization model proposed in [39] to account for admixture-LD while allowing for multiple generations and ancestries.

Although in our framework we use standard logistic gradient functions that were previously used to link geography and genetic variation, it is worth mentioning that such functions do not capture the whole variability observed in empirical data. To that extent, introducing more flexibility in these functions within the framework for admixture we described here are likely going to provide considerable improvements in accuracy with a tradeoff of computational time. We view this as a promising direction for future study. This is especially important for handling sequencing data, as rare variants rarely are fit well by the gradient functions (results not shown).

Another area for further developments is extending the framework to model background LD (correlations among variants on the same ancestral backgrounds). We found it necessary to modify the transition rates used in our inference by a multiplicative factor based on the level of LD pruning applied to the SNP list (Supp Table 2). Such LD adjustments have proved fruitful in improving localization accuracy for un-admixed individuals [3] and are likely to improve inference for admixed individuals as well. Although we leave this for future work, one potential approach would be perform inference within short windows (to account for the local structure of LD) and merge the information within each window into the overall likelihood.

We also note that we used a simplified model of ancestry switching along chromosomes that approximates the pedigree structure. In effect, our approach assumes a fixed effective time-scale of admixture, and ignores the structured transition matrices that are expected due to a fixed pedigree. Future work that explicitly considers the pedigree structure could allow one to address questions regarding the specific timing and configuration of admixed ancestries. For example, for a mixed individual with one Italian and three British grandparents, we could incorporate the specific inheritance pattern in the HMM transition rates. The question of whether the ancestors themselves were admixed could be investigated by assigning local ancestry followed by analyzing the length distribution of the ancestry blocks, and we leave this as future work.

For most of the results presented, we assumed that the number of ancestral locations is known. In practice, such information will often not be available. To address this we developed a procedure for inferring the number of ancestries using AIC. In Figure 3 we showed that while on average the correct number of ancestries will be inferred, in a high proportion of cases the inferred number of ancestries will be mistaken. This type of model selection problem is akin to estimating K in the admixture model of STRUCTURE/admixture [10, 1] and is typically challenging. In future work, pedigree-based models should lead to constraints in the possible observations that make the number of ancestors more straightforward to infer.

In this paper, we focus on the prediction of ancestral locations using an EM algorithm, which is a deterministic method to produce point estimates of the parameters of interest (geographic origins of ancestors) and missing data (the local ancestry of each allele copy). Alternative inference approaches can be taken-for example, the likelihoods we define in Equations (2) and (3) could be used in a Bayesian Markov Chain Monte Carlo (MCMC) approach method to sample from the posterior distribution of the spatial prediction[37]. In such an approach, an efficient starting point would be from the point estimate obtained via the EM algorithm, which could significantly expedite the convergence of the MCMC approach.

Our work provides a framework of predicting ancestral locations for admixed individuals which can be further improved. For example, due to the continuous nature of the approach, ancestral locations can be predicted to be outside of standard geographic boundaries (e.g. ocean). Future work could exploit this to improve on our framework by providing constrains to the optimization procedure or by a post-optimization adjustment (selecting the closest location that fits geographical boundaries). We leave a full investigation of such modifications of our framework as ongoing work.

A direct benefit of the proposed model is that it leads to efficient optimization procedures for tackling inference problems such as localization of ancestors in the genetic-geographic map. In this work we have presented such an algorithm based on the well known Expectation Maximization procedure that leverages the hidden Markov modeling of the admixture process joint with the gradient representation of genetic variation as function of geography.

## Methods

### Spatial modeling of allele frequency

Although our base method for explicit modeling genetic variation as function of geography has been described elsewhere [39, 3], we briefly present here the generative model. We view an individual’s alleles as a Bernoulli draw from an allele frequency that changes across the map and we parametrize the allele frequency function through a logistic gradient as function of position 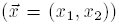 in the map. Formally, the probability of observing the reference allele in SNP *j* at position 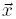 on the map 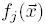, is defined as:

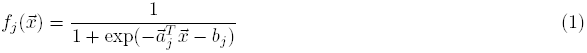

where 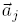 and *b*_*j*_ are parameters specific to SNP *j*. We estimate 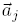 and *b*_*j*_ from data containing individuals with known homogeneous locations [39] and then use these coefficients in the inference of ancestries of mixed individuals.

Although easy to manipulate mathematically, the logistic functions we employ here clearly do not capture all genetic variation (for example, variants that have multiple modes or peaks in the allele frequency surface as may be typical of rare variants). However, these functions have been shown to capture general trends in common variant frequencies sufficiently well to produce highly accurate spatial assignment in individuals with non-mixed ancestry [39]. We hypothesize that such simple-to-manipulate functions are sufficient for accurate localization of recent ancestors in individuals with mixed sub-continental ancestries.

### Haploid data with admixed ancestry

#### Spatial model for admixed haploid data

For simplicity, we start by introducing the model for haploid data and extend it to genotype data in the next section. Denote by *h* = (*h*_1_, …, *h*_*L*_) the multi-site haplotype of an admixed haplotype, where *L* is the number of SNPs typed across the genome and *h*_*i*_ ∈ {0, 1} encodes the number of reference alleles at SNP *i*. Due to the admixture process, the haplotype can be viewed as a mosaic of segments coming from ancestors from multiple locations on the map. We define variables *Z* = (*z*_1_, …, *z*_*L*_) as indicators for an allele coming from ancestry location *j* (*z*_*i*_ = *j* if allele at locus *i* is from *j*-th ancestry location) and write the likelihood of the haplotype data as function of ancestry locations *X*. The likelihood for a given admixed haplotype data having *M* ancestry locations *X* = (*x*_1_, …, *x*_*M*_) where each ancestry contributes proportionally with π = (*π*_1_, …, *π*_*M*_) is defined as:

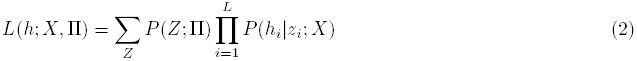

The hidden variable *Z* encodes the mosaic structure of the admixed haplotype (i.e. inheritance within the past generations for recent admixture, admixture-LD) and can be modeled using a Markov chain as follows:

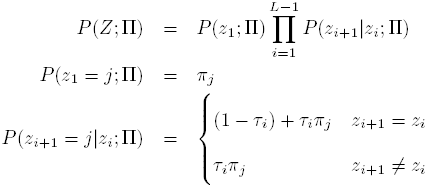

where the parameters τ = {τ_1_, …, τ_*L*−1_} stand for the recombination probability (within the past *g* generations) between each two neighbor loci. The alleles present at a site *i* on a haplotype is modeled as a Bernoulli variable with a success probability given by the allele frequency *f*_*i*_(*x*_*z*_*i*__) as follows:

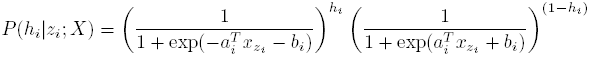

An illustration of the model is given in Figure 1. We note that our model makes the assumptions of independence of alleles conditional on local ancestry (no modeling of background LD nor deviations from Hardy-Weinberg proportions).

#### Spatial ancestry inference for haploid data

Under the generative model above, spatial ancestry inference is reduced to inferring the *M* ancestral locations given data for an admixed haplotype, followed by posterior decoding in the HMM to obtain locus-specific predictions. This can be achieved by maximizing the likelihood function (2) with respect to *X*. By treating *X* as parameters and *Z*, π as hidden variables, this maximization falls within the procedure of the standard Expectation Maximization (EM) algorithm [8]:

*E step:* The expectation step is similar to the forward-backward algorithm for Hidden Markov Models (HMM) which calculates the posterior probability of hidden variables *Z* given current estimation of ancestral locations *X*^(*t*)^ and ancestral proportion π^(*t*)^:

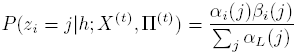

where *α*/*β* are standard forward/backward HMM functions and can be efficiently calculated (see Supplementary Note).

*M step:* The maximization step alternatively optimizes the Q functions in *X* and in π [8]. The first can be derived as

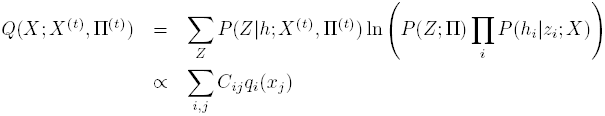

where *C*_*ij*_ denotes the posterior *P* (*z*_*i*_ = *j*|*h*, *X*^(*t*)^, π ^(*t*)^) from the *E step*, and the shorthand *q*_*i*_(*x*_*j*_) is defined as:

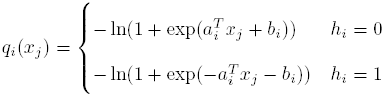

The Q function in π can be derived as follows

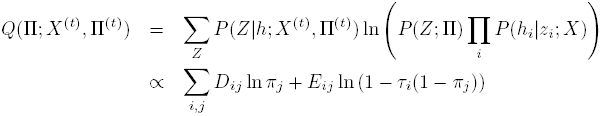

where *D*_*ij*_ and *E*_*ij*_ denote constants calculated from the posterior *P* (*z*_*i*_ = *j*|*h*, *X*^(*t*)^, π^(*t*)^) in the *E step*.

We perform the maximization by taking advantage of the convex properties of the equation and using analytical forms for the Hessian of the function. The complete derivations are given in Supplementary Note.

#### Locus-specific spatial ancestral inference for haploid data

Having obtained the maximum likelihood geographical locations *X*^*^, we compute the posterior probability for *Z*, which leads to a locus specific assignment of ancestry at each allele in the genome. The most probable local ancestral locations are found by maximizing

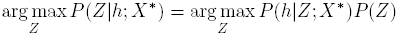

which can be efficiently solved by the Viterbi algorithm [36]. In order to compute a posterior probability of each locus-specific ancestry, we employ the forward-backward algorithm (see Supplementary Note).

### Diploid data with admixed ancestry

#### Spatial model for admixed diploid data

We next extend the haploid model to genotypes by considering *M* paternal ancestry locations *X* = (*x*_1_, …, *x*_*M*_) with proportions П = (*π*_1_, …, *π* _*M*_) and *N* maternal ancestry locations *Y* = (*y*_1_, …, *y*_*N*_) with proportions Ω = (*ω*_1_, …, *ω*_*N*_). Denote by *g* = (*g*_1_, …, *g*_*L*_) the multi-site genotype of an admixed genotype, where *L* is the number of SNPs typed across the genome and *g*_*i*_ ∈ {0, 1, 2} encodes the number of reference alleles at SNP *i*. Then the likelihood becomes:

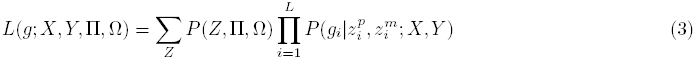

The variables *Z*^*p*^ and *Z*^*m*^ now encode the ancestry status of the paternal (maternal) alleles (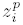 = *j* denotes that the paternal allele at locus *i* is from *j*-th paternal ancestry), and can be modeled through the same Markovian process as:

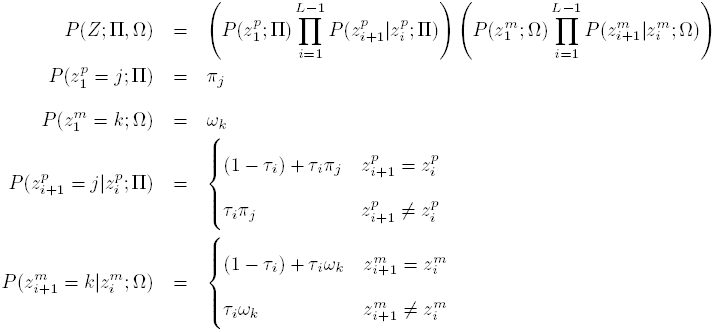

Given the origin of alleles, the likelihood of the admixed individual genotype is modeled as two Bernoulli draws:

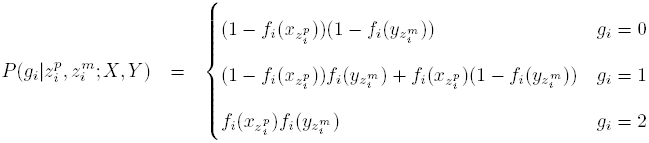

The function *f*_*i*_ is the allele frequency function in logistic form (1). The probability *P* (*Z*) models the recombination events in paternal and maternal ancestries, and the probability 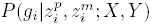 models the probability of generating the genotype from two ancestral geographical locations.

#### Spatial ancestry inference for diploid data

We would like to infer *M* + *N* ancestral locations for a given mixed individual genotype. This can be achieved by maximizing the likelihood function (3) with respect to *X* and *Y*, which, analogous to the haploid case, can be performed using the EM algorithm [8]:

*E step:* In short, the expectation step is similar with forward-backward algorithm in HMM, which calculates the posterior probability of hidden variables *Z* given current estimation of ancestral locations *X*^(*t*)^ and *Y*^(*t*)^.

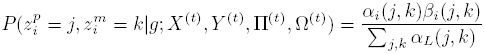

where *α* and *β* can be calculated recursively using a procedure similar to the forward-backward algorithm for HMMs.

*M step:* The maximization step alternatively optimizes the Q functions in *X*, *Y* π and Ω. The first can be derived as follows

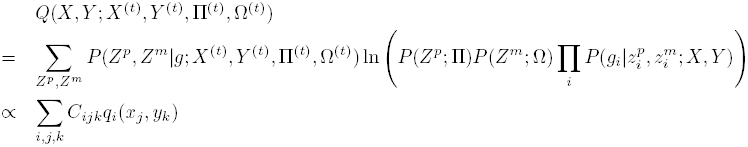

where *C*_*ijk*_ denotes the posterior 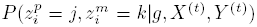 computed from *E step*, and the shorthand *q*_*i*_(*x*_*j*_, *y*_*k*_) is defined as:

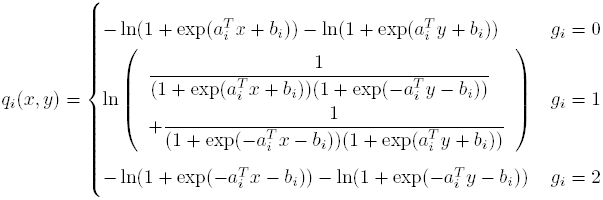

The Q function in π can be derived as

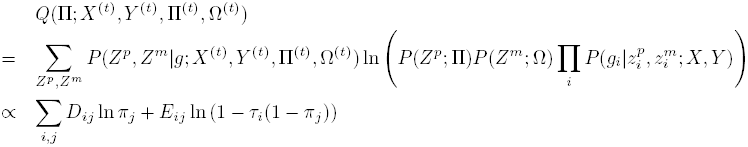

where *D*_*ij*_ and *E*_*ij*_ denote constants calculated from the posterior 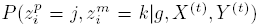 in the *E step*. We omit the Q function in Ω here as it is very similar with the above function.

As in the haploid case, we leverage the convexity of the function and analytical forms for the Hessian to efficiently optimize the Q function. The complete derivations and optimization details are given in the Supplementary Note.

#### Locus-specific spatial ancestral inference for diploid data

Having obtained the maximum likelihood geographical locations *X\** and *Y\** for each ancestry, we can compute the posterior probability for *Z*^*p*^ and *Z*^*m*^, which leads to the spatial local ancestry inference. The most probable local ancestral states is obtained by maximizing

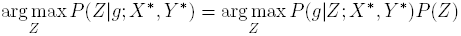

which can be efficiently solved by the Viterbi algorithm [36]. The posterior of local ancestries for each allele can be obtained using a forward-backward algorithm following the *E step* in the algorithm (see Supplementary Note).

#### Homogeneous paternal and maternal ancestries

In notations above we derived the general solution that allows for paternal and maternal ancestries to be different from each other, which is suitable for applications of inference of parental locations or grandparent locations. A simplifying case is when maternal and paternal ancestries are homogeneous; i.e. the paternal haplotype and maternal haplotype are from the same set of ancestral populations. We allow for this case by setting *M* = *N* and enforce a constraint *x*_*j*_ = *y*_*j*_ in the M step.

### POPRES data set

We applied our methods to a data set collected from European populations, which was assembled and genotyped as part of the larger POPRES project [19] and accessed via dbGAP accession number phs000145.v4.p2. A total of 3192 European individuals were genotyped at 500, 568 loci using the Affymetrix 500K SNP chip. The same stringency criteria as in [20] were applied to create the training data. We removed SNPs with low-quality scores and high missingness[20]. We filtered individuals to avoid sampling individuals from outside of Europe, to create more even sample sizes across Europe, and to remove individuals whose self-reported data have grandparents with different origins. We note that this is the same data set used in [20, 3]. For the remaining individuals who have observed grandparental data, we use that origin for the individual. Otherwise, we use the individual-level self-reported country of birth. As a result, we infer logistic gradients starting from genotype data from 447, 245 autosomal loci in 1, 385 individuals from 36 populations. 77.4% of SNPs are common SNPs (allele frequency > 0.05), and the remanining 22.6% have low frequencies (< 0.05). For testing, we identified an additional 470 individuals from the POPRES data that have self-reported grand-parental ancestry from 2 or more countries in Europe. A summary of homogeneous ancestry individuals used in estimating logistic gradients (1,385) and with sub-continental European admixed ancestry (470) are given in Table 1.

### Accounting for background Linkage Disequilibrium

Although our approach models adixture LD, it assumes that markers are independent conditional on local ancestry (no background LD). Preliminary results (not shown) that used transition rates based on the assumed number of generations and recombination rate (i.e. similar to simulations, (*g −* 1)*φ*(*d*_*i*+1_ − *d*_*i*_) where *d*_*i*_’s are the locations of each SNPs) attained increased number of short ancestry windows which leaded to decreased accuracy. This effect is likely due to lack of modeling of background LD in the model. To remove short ancestry windows (likely spurious, induced by residual LD) we first performed LD pruning at a level of *r*2 < 0.2 (72,418 SNPs retained) followed by adjustment of the transition rates in our model by a factor of 10^−2^. Results at different LD pruning levels and adjustment factors are reported in Supplementary Tables 1, 2.

### Simulation setup

We use BEAGLE to phase the POPRES data then simulate offspring admixed individuals by modeling recombinations within the last couple of generations. The recombination probability between each SNPs is approximated as (*g −* 1)*φ*(*d*_*i*+1_ − *d*_*i*_) where *d*_*i*_’s are the locations of each SNPs in bp, *g* denotes the number of generations and *φ* is the probability of one recombination per generation per base-pair [21]. For the recombination map, we assumed a flat recombination rate of *φ* = 10^−8^ per base-pair. For given number of *M* paternal ancestries and *N* maternal ancestries, we randomly select from the POPRES data a set of *M* + *N* individuals, each of which has 4 grandparents from the same locations and randomly select one haplotype from each individual. We simulate the admixed haplotypes independently for the maternal and paternal haplotypes using the standard Poission process of admixture block distribution [28]. If specified as homogeneous paternal and maternal ancestries, we pick *M* instead of *M* + *N* ancestries, and use the same *M* ancestries for both paternal and maternal haplotype simulation.

For the SPAMIX haploid model, the simulated haplotypes are used as input directly. Also, we always use the correct number of ancestries *M* or *N* as input. For the SPAMIX diploid model, the combined genotype from two simulated paternal and maternal haplotypes are used as input. To avoid testing bias, we estimate the allele frequency logistic gradients each time using the POPRES individuals with the *M* + *N* simulation ancestors excluded from the training set. We do not optimize over the ancestry proportions but provide them as input to SPAMIX.

We use several metrics to assess performance of SPAMIX in simulations and real data. For the ancestral location prediction, we evaluate the results by computing the average geographical distance between predicted locations and true locations in simulations (*prediction error*). To account for the distance among ancestries we also compute the *relative prediction error*, defined as the ancestral location prediction error divided by the distance between the true ancestry locations used in simulations. Note that we use as the “true” ancestral locations for the admixed individual the set of country centers from the *M* + *N* ancestries.

For locus-specific inference, we propose two different metrics. The first one is the *local ancestry prediction error*, which is the average distance between predicted location and true location at each locus. The second metric we use is the *local ancestry prediction accuracy*, defined as the percentage of loci across the genome with correct assignment of ancestry. To account for the ambiguity in matching the true to inferred ancestries, we permute the inferred ancestries to find the closest match in terms of inferred location to true location.

**Figure 5:**
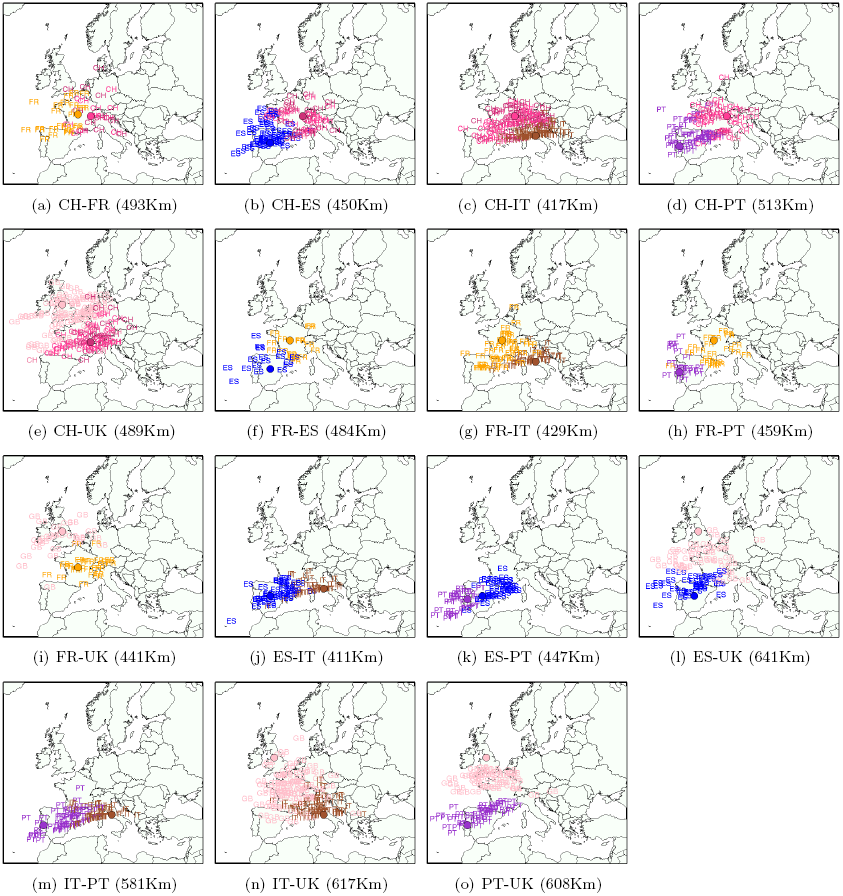
Ancestral location prediction error in simulations of European individuals with ancestry from two locations in Europe, stratified by the country of origin of each location (the country of origin is displayed in different colors). The assumed true locations are displayed by shaded circles. Results in parenthesis denote the average ancestral location prediction error across all simulations. In each simulation the reference data (used to estimate logistic gradients) is disjoint from data used to simulate admixed genomes (see Methods). The admixed genome is simulated as 4 generations ago, and SPAMIX diploid model is used for the inference. The number of simulated pairs can be found in Figure S4.

**Figure 6:**
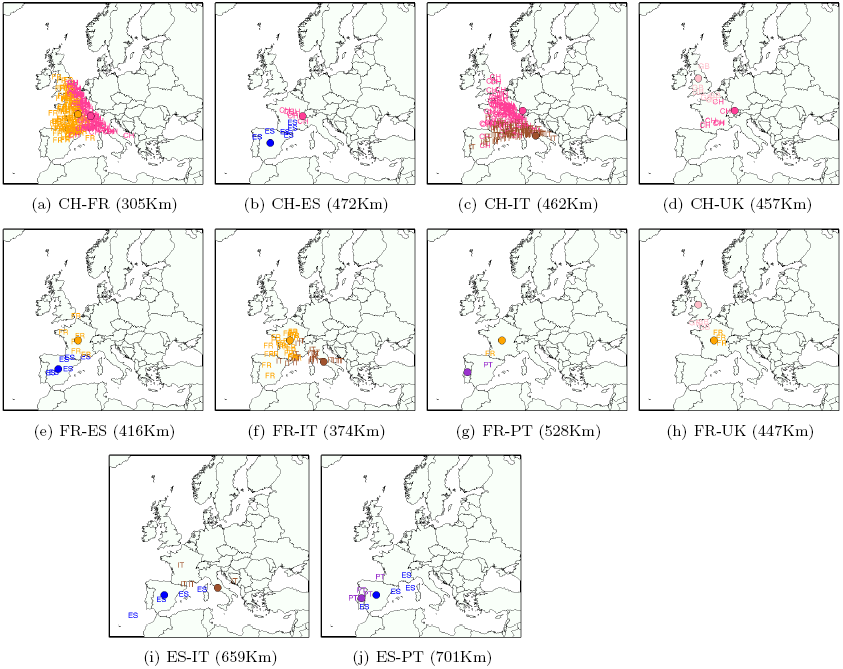
Ancestral location prediction error in real POPRES admixed individuals, stratified by the country of origin of each location. Letters are the inferred locations, and the shaded circles are the assumed true locations.

